## Supplemental Figures for "A specific role for Importin-5 and NASP in the import and nuclear hand-off of monomeric H3"

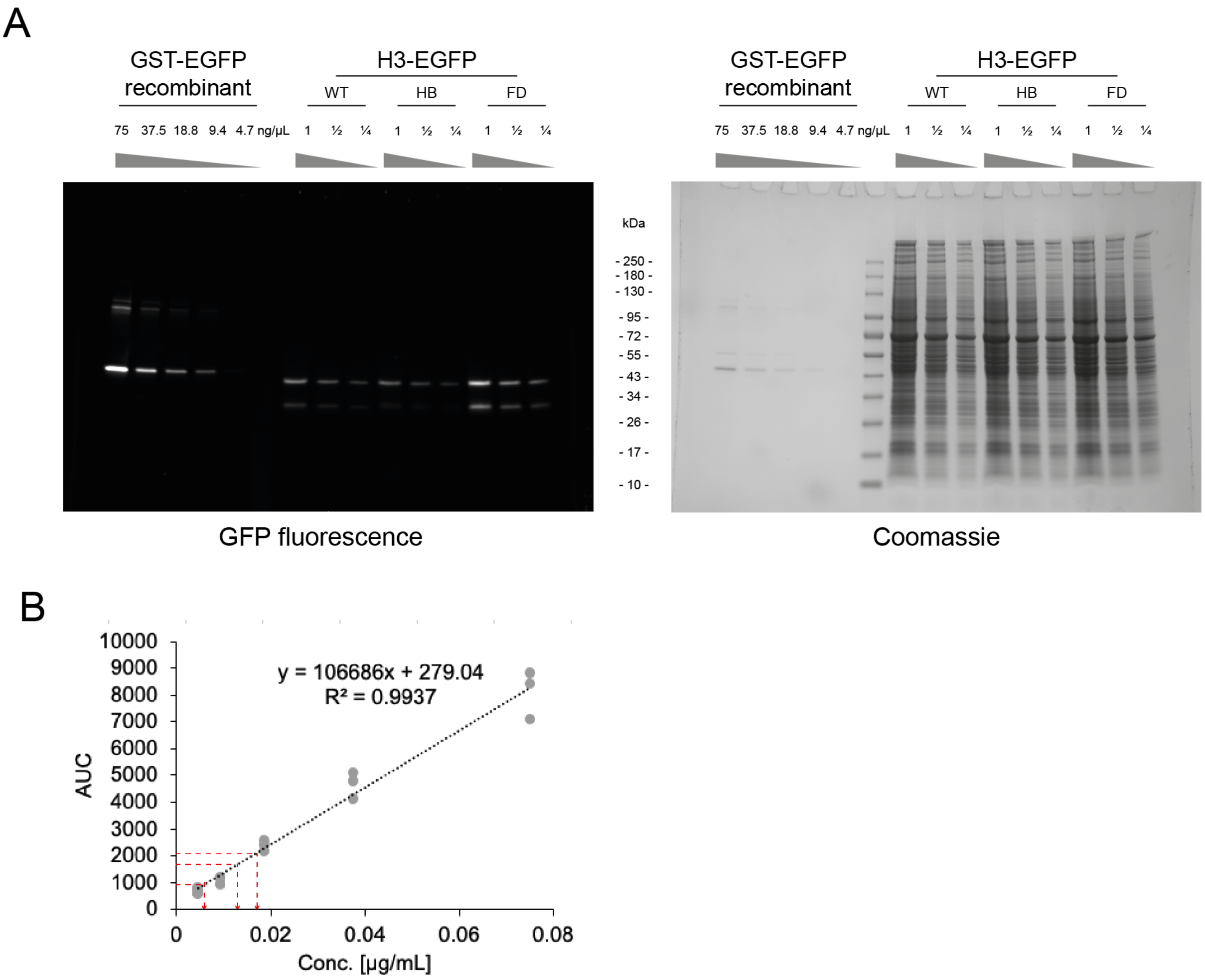


**Figure S1**. Related to Fig 2. Calibration for precise input quantification.
A: Residual fluorescence activity of GFP constructs on a semi-denaturing gel before staining (left) and after Coomassie staining (right). B: Area under the curve (AUC) analysis of residual GFP fluorescent bands, for GST-GFP calibration curve and sample dilutions (red line).

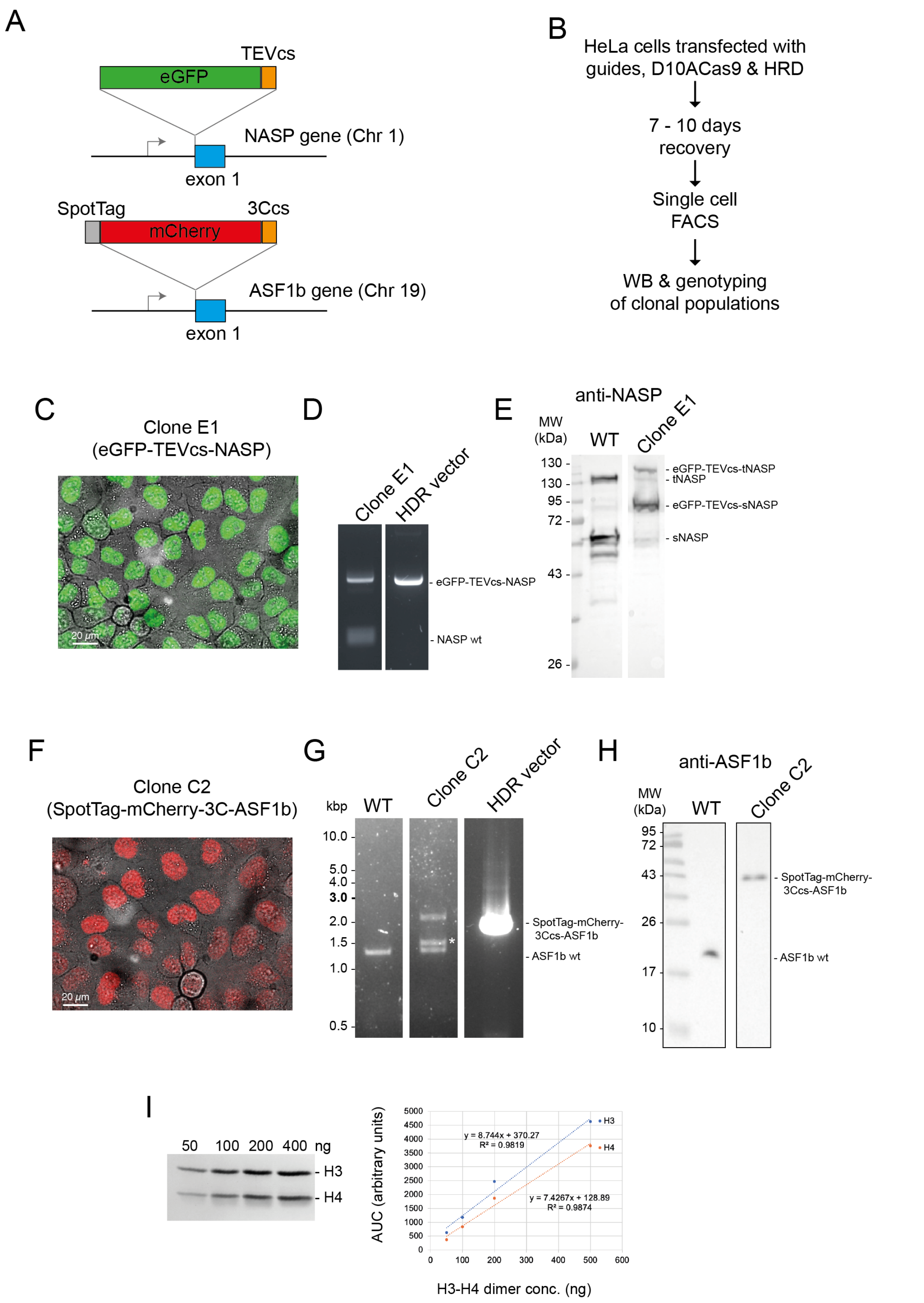


**Figure S2**. Related to Fig 3.
A: Experimental design for CRISPR Knock-in cell line selection, western blot (WB) and PCR validation of the clonal lines. B: Confocal microscopy for selected CRISPR Knock-in cell lines EGFP-NASP and mCherry- ASF1B, demonstrate nuclear localisation, even expression and complete penetration. C: Serial dilutions for H3-H4 quantification. 50–400 ng of H3-H4 dimer was separated by SDS–PAGE and stained with Coomassie as previously described (Apta-smith et al., 2018).

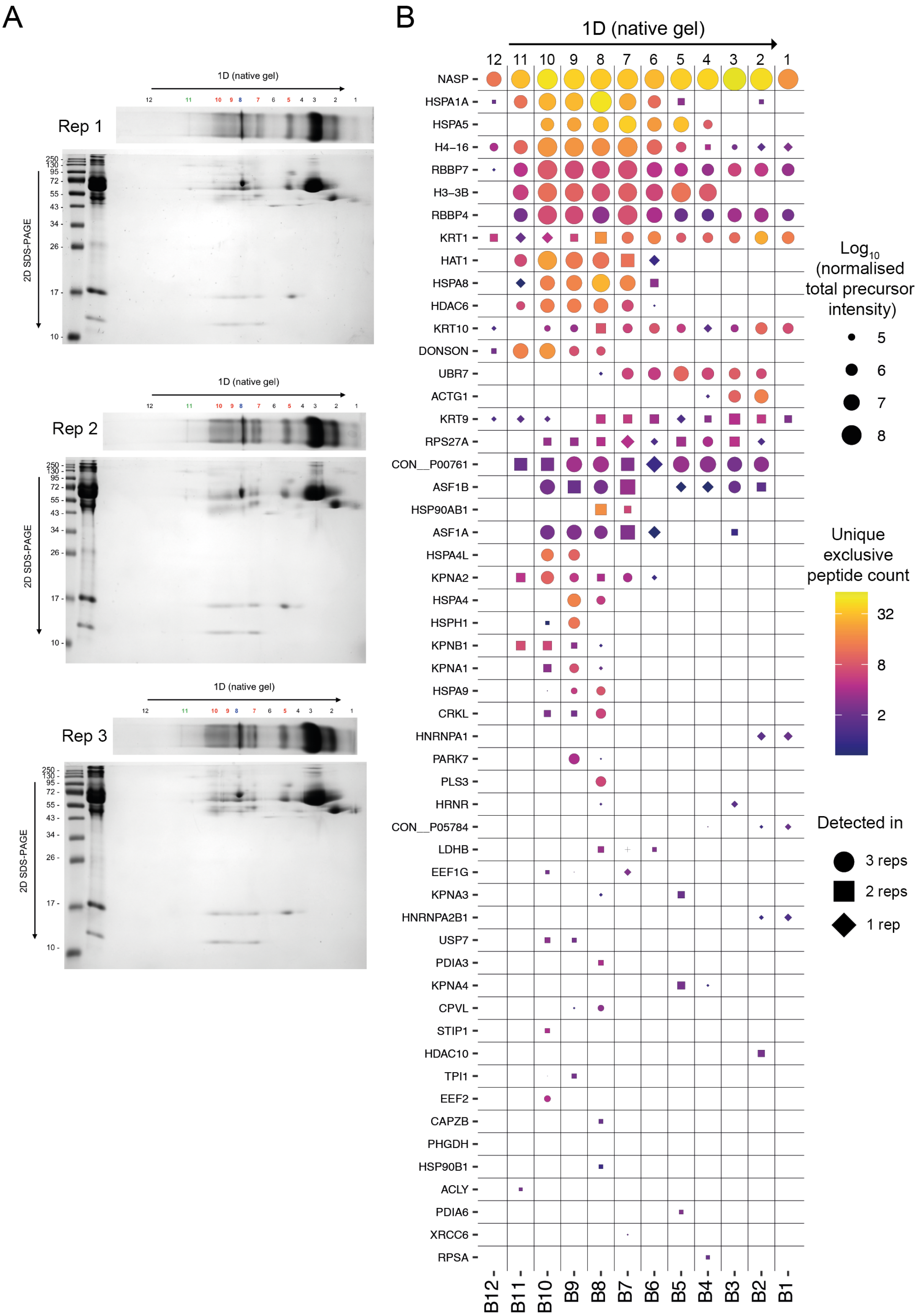


**Figure S3.** Related to Fig 3.
A: Native gel (1D) and SDS-PAGE (2D) for 3 independent experiments (performed with different batches of cell transfected on different days, processed subsequently 36 hrs after transfection). B: Mass-spectrometry identification of NASP-interacting factors from native gel bands (cut as labelled in A).

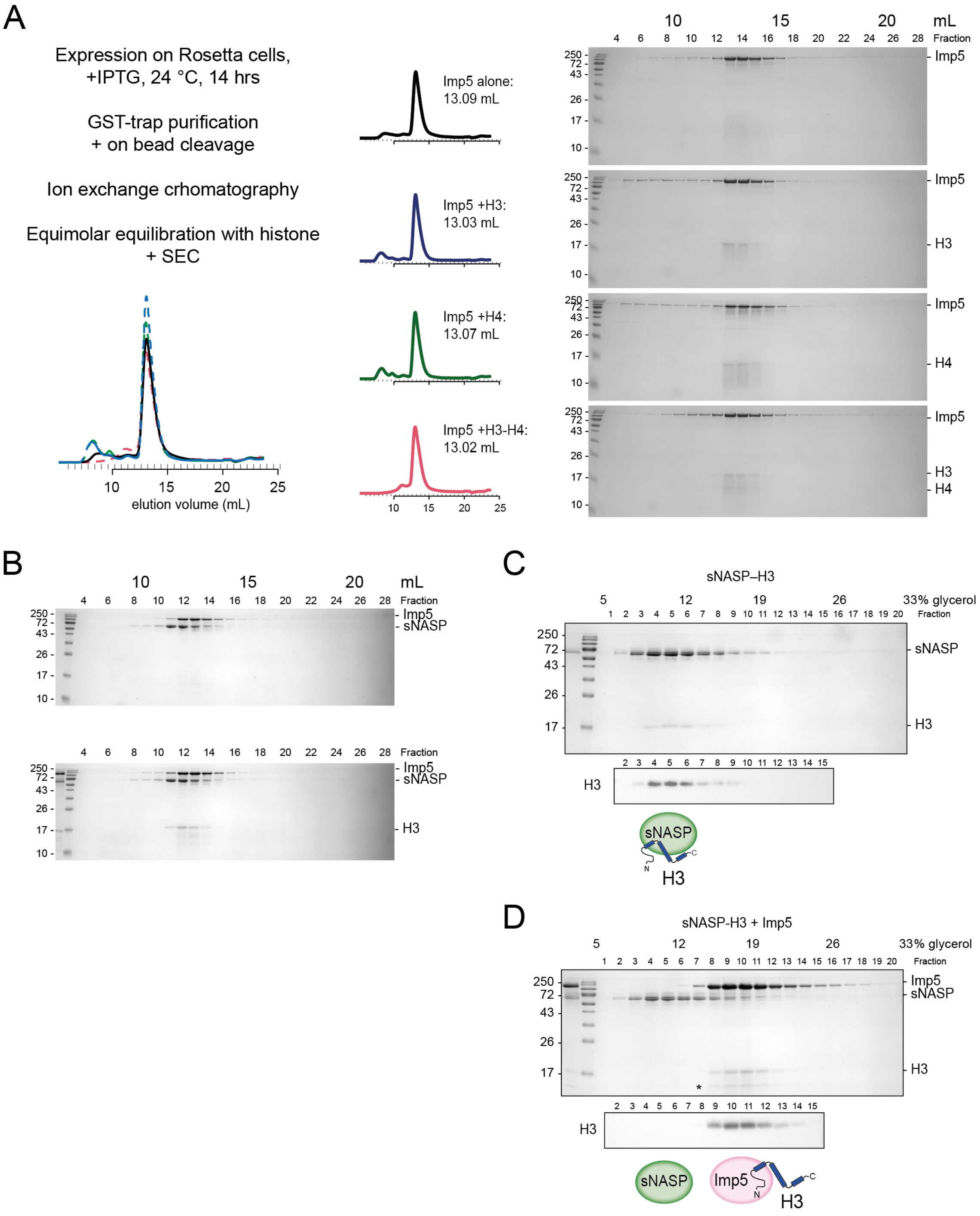


**Figure S4.** Related to Fig 4.
A: Experimental design for Imp5 purification and complex reconstitution, SEC$_{280}$ trace (on Superdex Increase 200), peak volumes (mL) and Coomassie-stained gels. B: SEC for sNASP + Imp5 does not resolve these proteins (i.e. elution peaks at ~12.6 and 13 mL respectively). C: 5-40\% glycerol gradient ultracentrifugation (240,000 g, 4 °C, 12 hours) sNASP-H3. D: Imp5 outcompetes pre-formed sNASP-H3 and displaces H3. (equimolar mix).

A

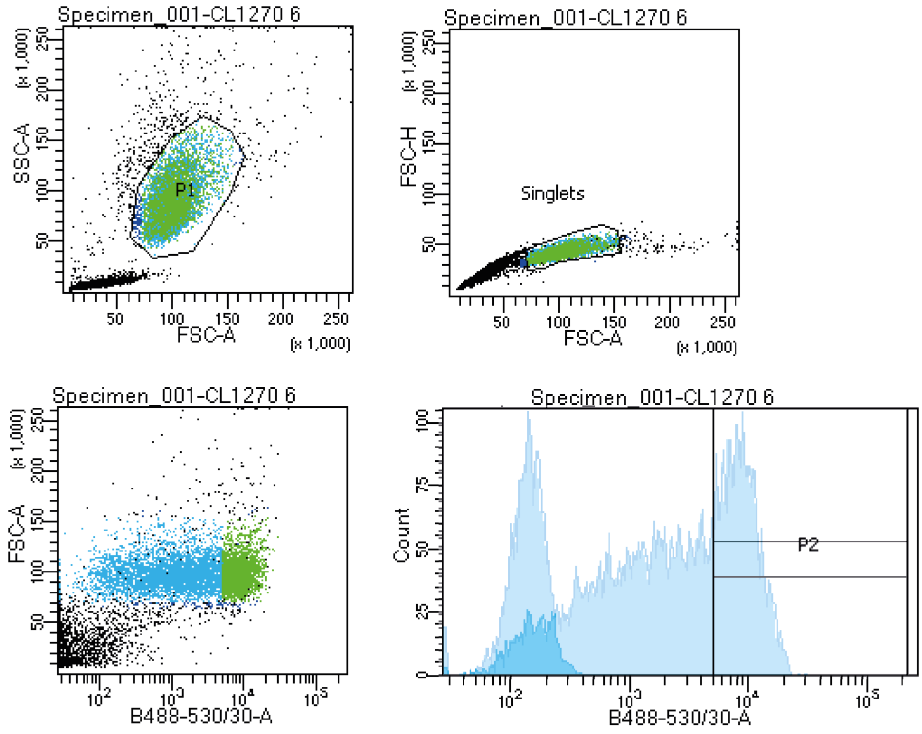


Tube: CL1270 6

| Population | #Events | %Parent | %Total |
|---|---|---|---|
| All Events | 10,000 | #### | 100.0 |
| P1 | 6,827 | 68.3 | 68.3 |
| Singlets | 6,731 | 98.6 | 67.3 |
| P2 | 2,352 | 34.9 | 23.5 |

B

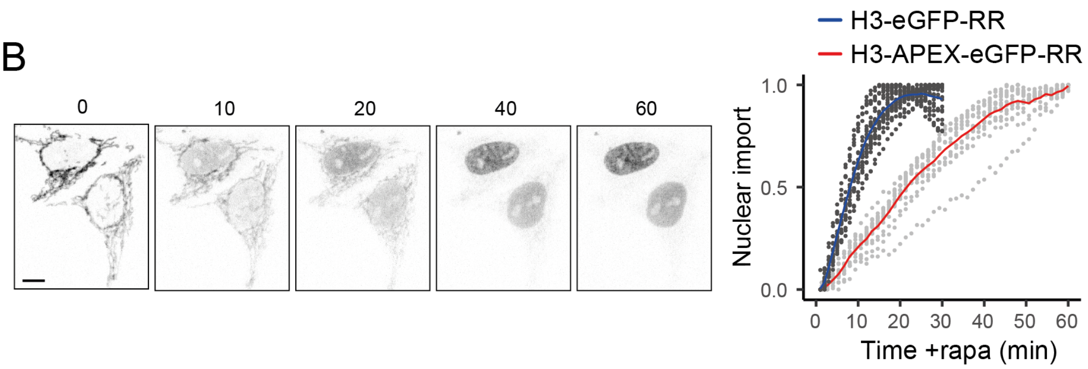


**Figure S5**. Related to Fig 5. A
A: FACS data for FRT -H3-APEX2-EGFP-RR cells induced with doxycycline for 12 hours, compared to non-induced cells (faded) in blue laser histogram (bottom right graph). B: Confocal microscopy time course of doxycycline-induced FRT-H3-APEX2-EGFP-RR and nuclear enrichment analysis (red) compared to H3-eGFP-RAPID release (RR) in blue. Note that the addition of the APEX2 tag delays the cleavage kinetics and nuclear localisation.

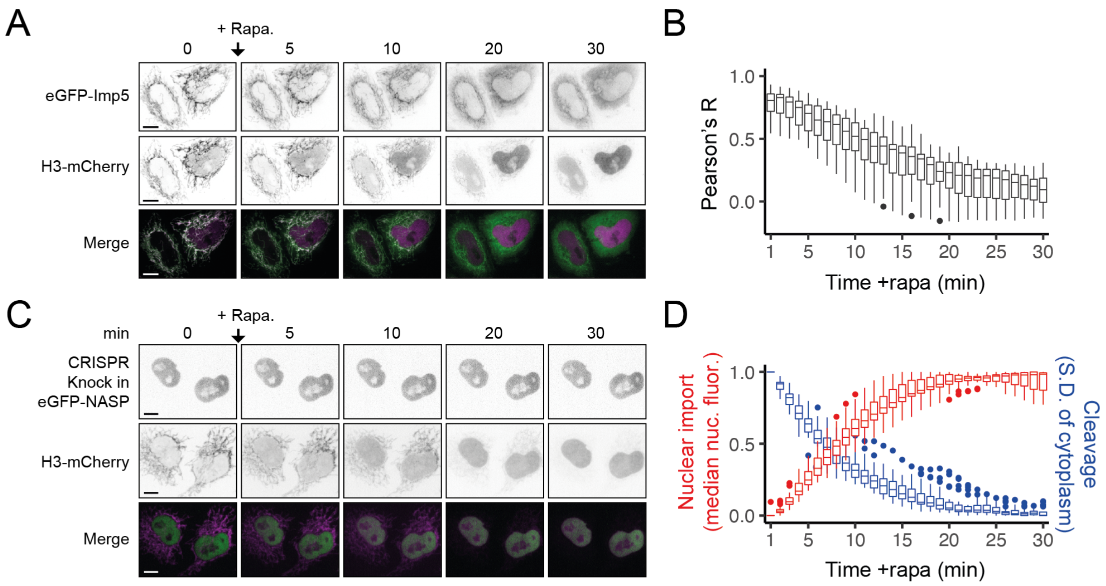


**Figure S6.** Related to Fig 6.
A: Confocal microscopy time course after the addition of rapamycin (RAPID release H3-mCherry-RR) in cells transiently expressing eGFP-Imp5. Bar = 10 µm. B: Pearson's correlation coefficient (R) analysis (as in A. n = 20 cells). C: Confocal microscopy time course after the addition of rapamycin (RAPID release H3-eGFP-RR) in HeLa cells with eGFP-NASP endogenously labelled (CRISPR knock-in). Bar = 10 µm. D nuclear enrichment analysis compared to cleavage rate (inferred from S.D of cytoplasm).
