## Supporting Material for "A specific role for Importin-5 and NASP in the import and nuclear hand-off of monomeric H3"

Fig 1E

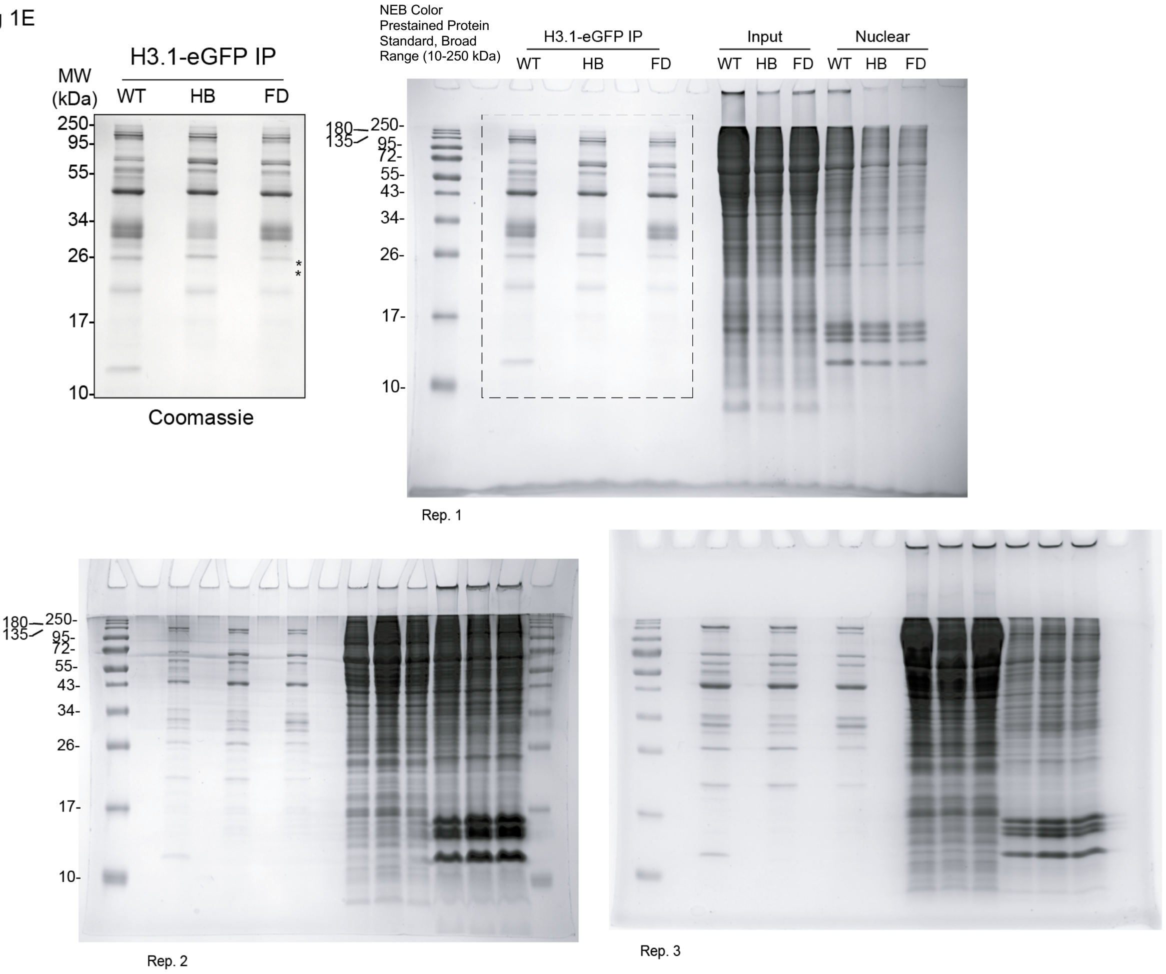

H3.1-eGFP IP
MW (kDa)
WT HB FD
250- 95- 55- 34- 26- 17- 10-
Coomassie
NEB Color Prestained Protein Standard, Broad Range (10-250 kDa)
H3.1-eGFP IP
Input
Nuclear
WT HB FD WT HB FD WT HB FD
180 135 250- 95- 72- 55- 43- 34- 26- 17- 10-
Rep. 1
Rep. 2
Rep. 3


Fig 1F

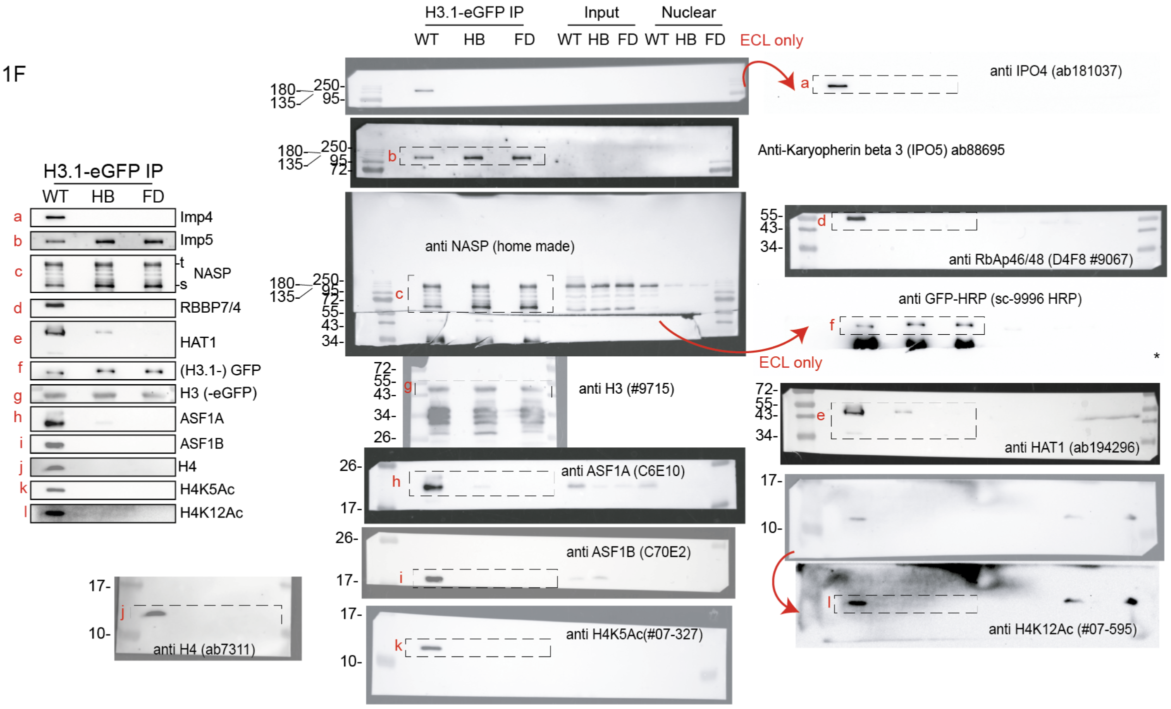

H3.1-eGFP IP
WT HB FD
a Imp4
b Imp5
c NASP t s
d RBBP7/4
e HAT1
f (H3.1-) GFP
g H3 (-eGFP)
h ASF1A
i ASF1B
j H4
k H4K5Ac
l H4K12Ac
anti H4 (ab7311)
H3.1-eGFP IP
Input
Nuclear
WT HB FD WT HB FD WT HB FD
ECL only
anti IPO4 (ab181037)
Anti-Karyopherin beta 3 (IPO5) ab88695
anti NASP (home made)
anti RbAp46/48 (D4F8 #9067)
anti GFP-HRP (sc-9996 HRP)
ECL only
anti H3 (#9715)
anti HAT1 (ab194296)
anti ASF1A (C6E10)
anti ASF1B (C70E2)
anti H4K5Ac(#07-327)
anti H4K12Ac (#07-595)

Fig 1E

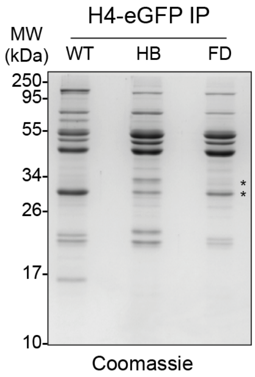


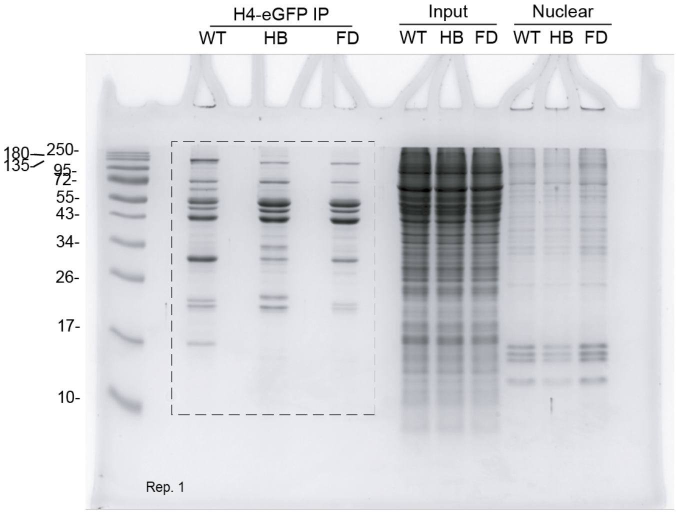


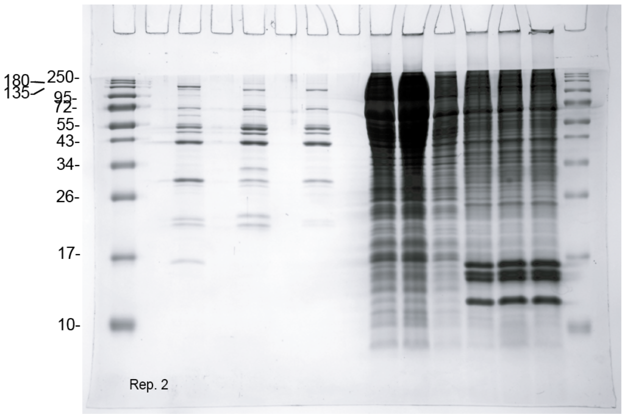


Rep. 3 Coomassie staining unavailable

Fig 1F

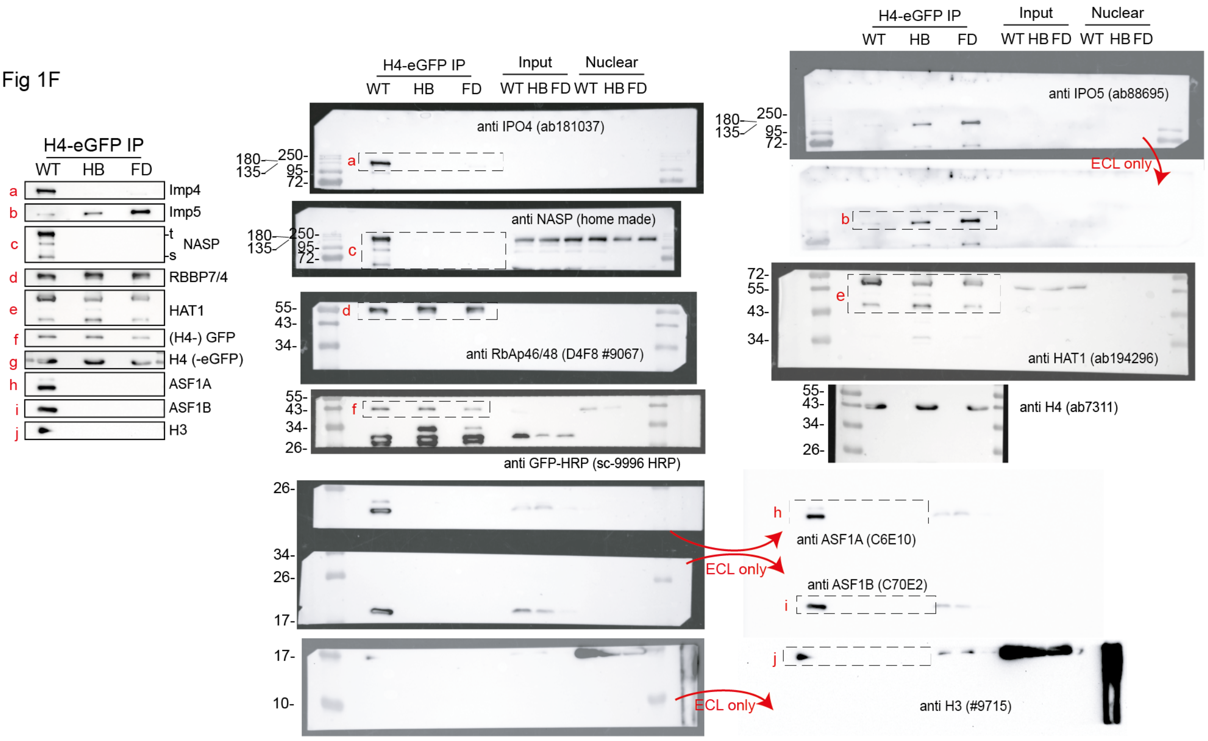

Fig 3B

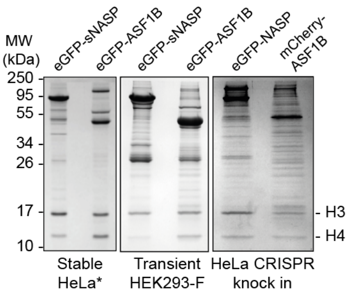


HEK293-F transient expression

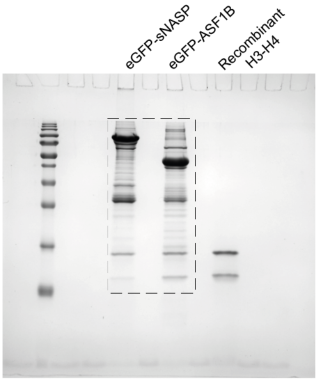


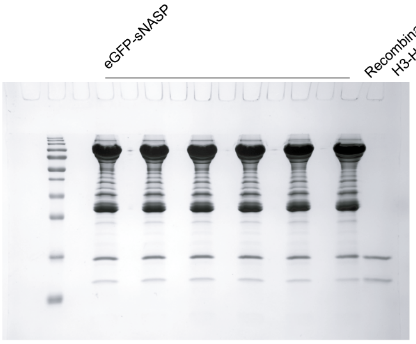


HeLa CRISPR knock-in endogenous expression

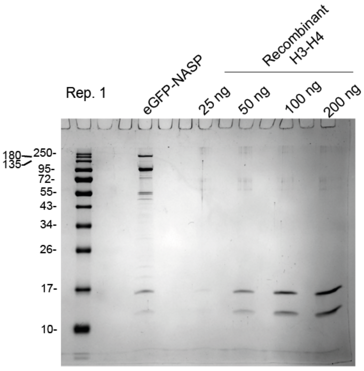


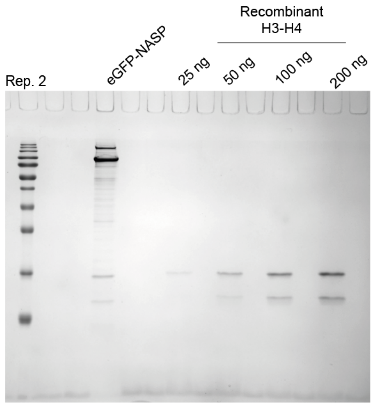


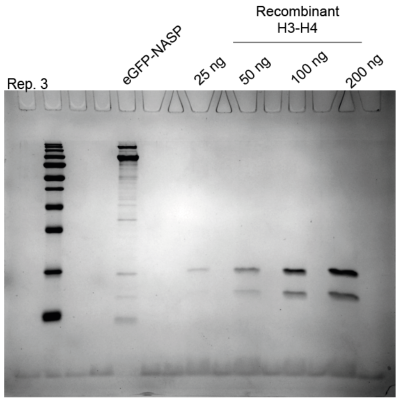


HeLa CRISPR knock-in endogenous expression:
eGFP-NASP vs.
mCherry-ASF1

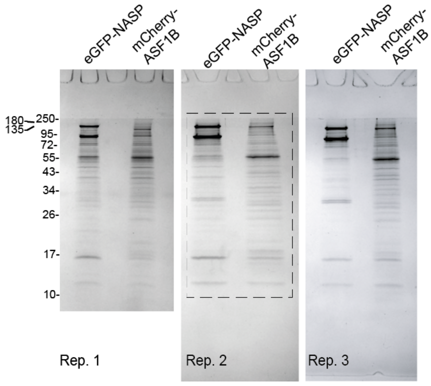

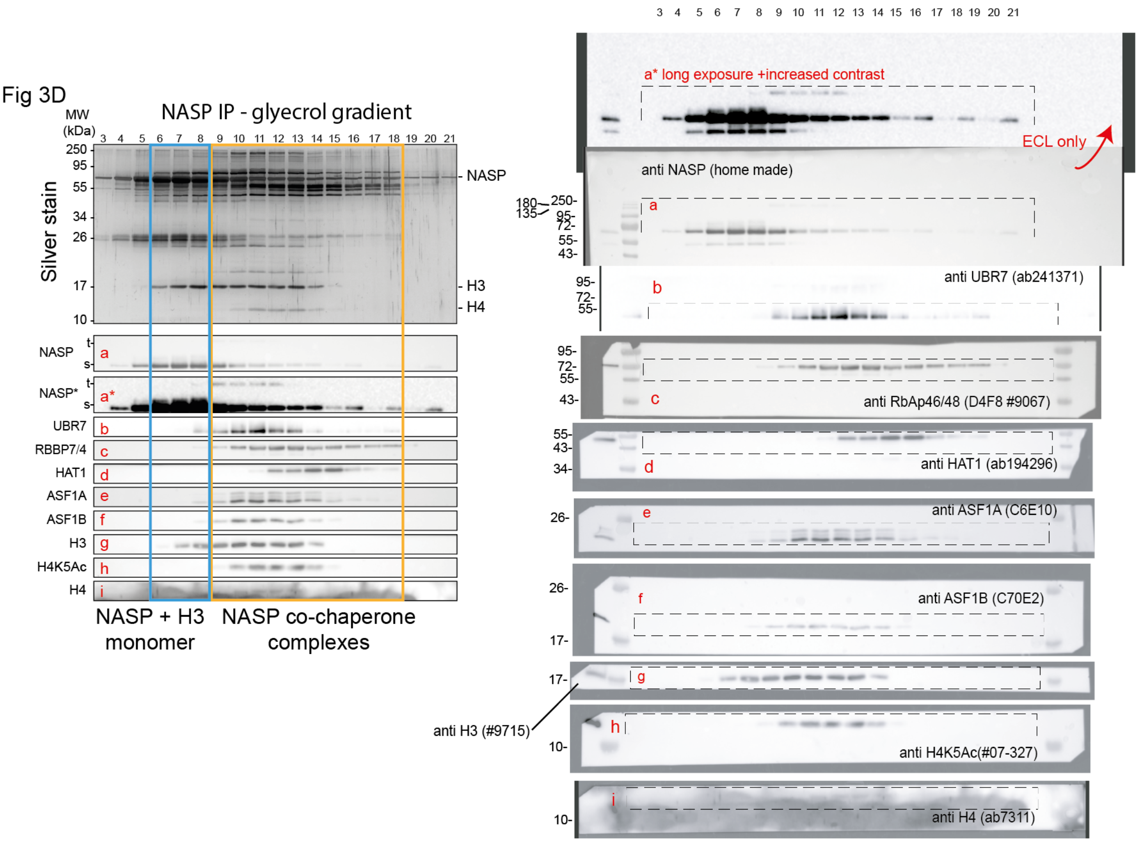


For replicates of the stained native gels (send for mass spectrometry) and 2D gels see Supp. Fig. 3

To avoid excessive background, anti NASP primary antibody was incubated with anti Rabbit alexa 561 conguated (unlike the remaining blots in this work, where the secondary antibody was HRP congugated)

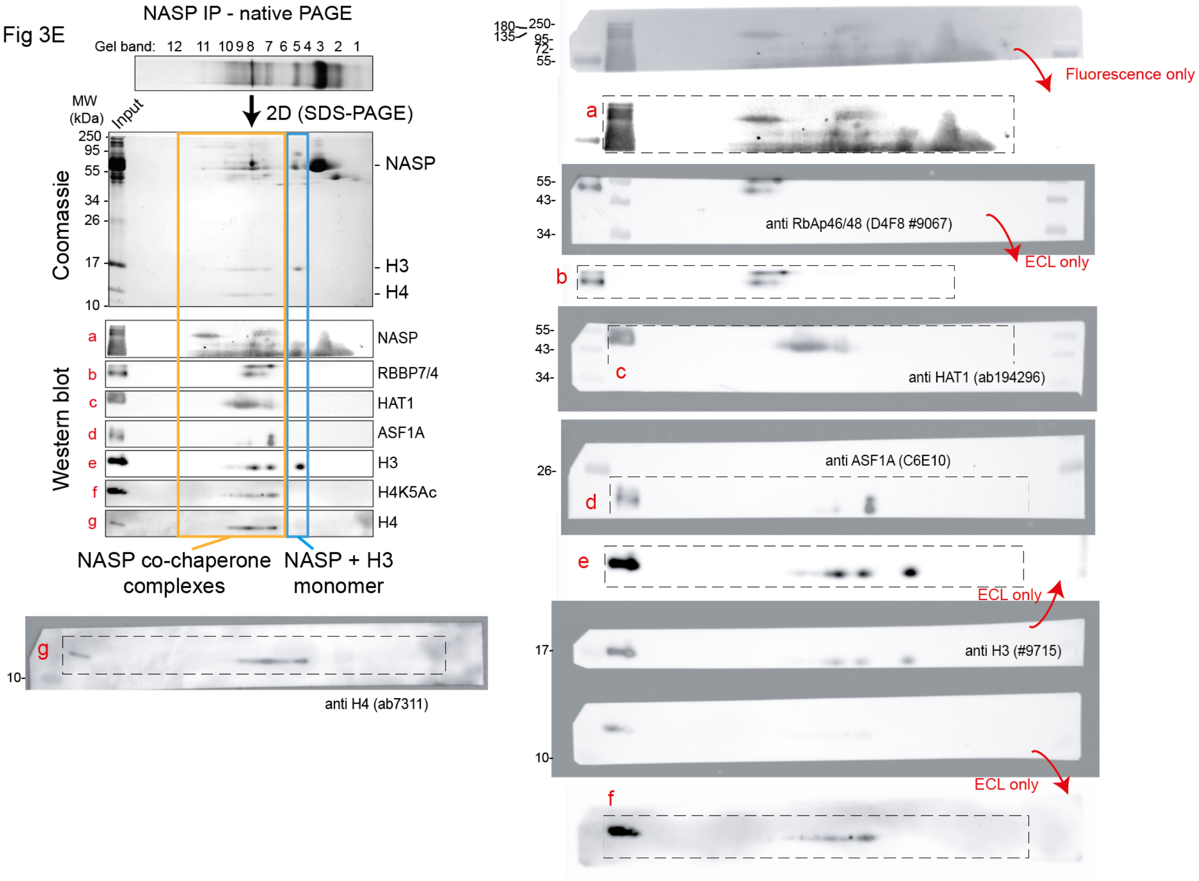

Fig 4A

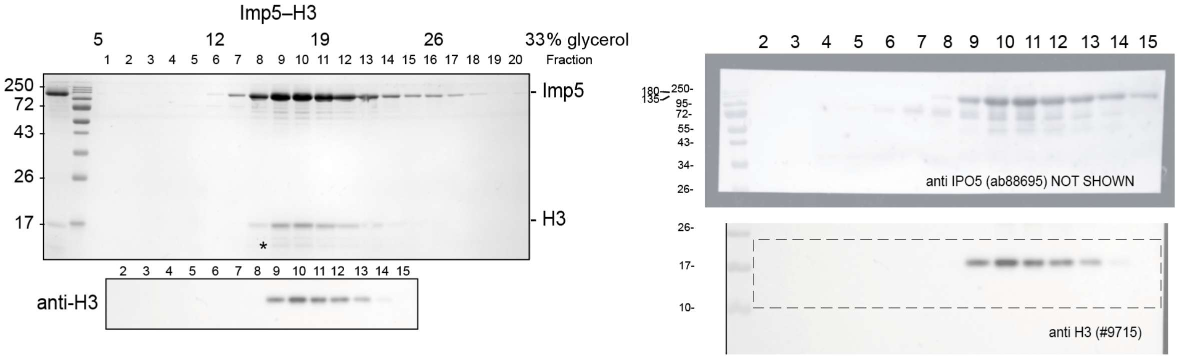


Fig 4B

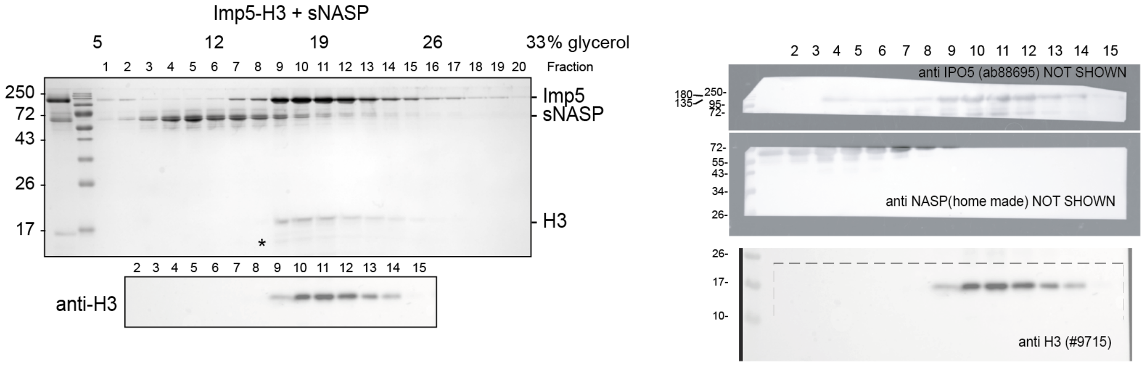


Fig 4C

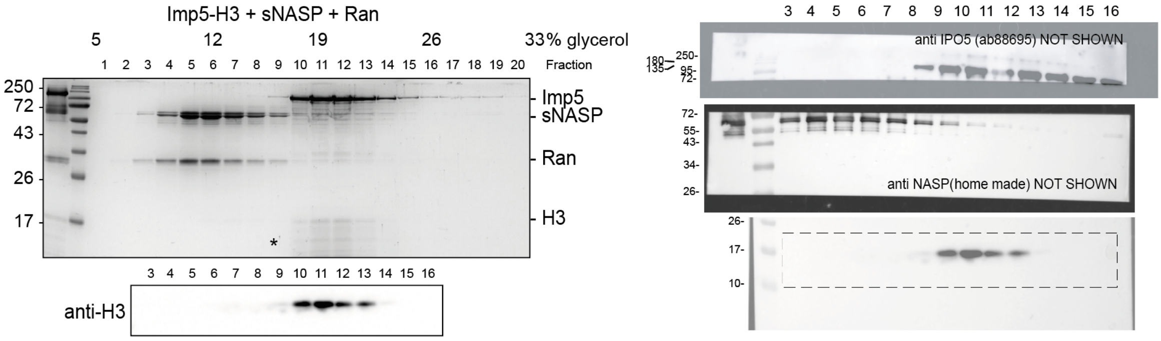


Fig 4D

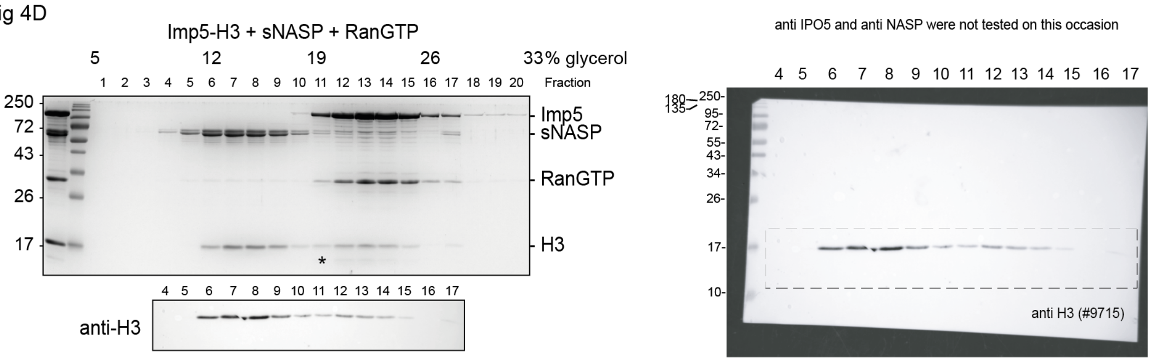

Fig 6 B

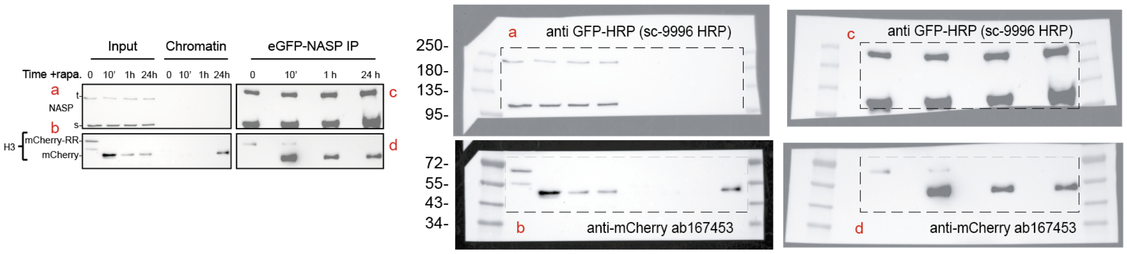


Rep. 2

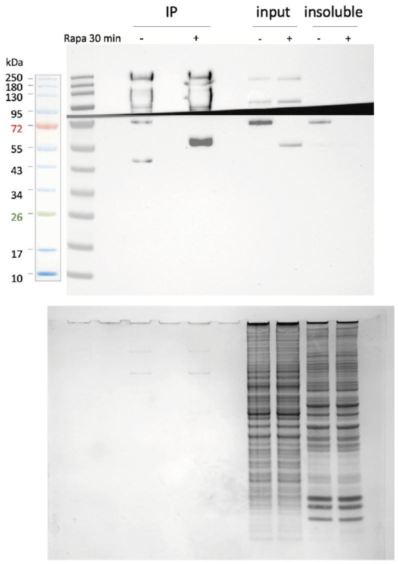


Rep. 3

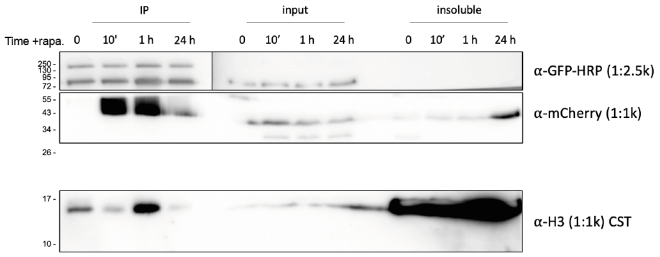
