## Supplementary material for "A specific role for Importin-5 and NASP in the import and nuclear hand-off of monomeric H3": Table S1

### Table S1. Plasmids used in this study.

| Plasmid name | Construct | Backbone | Origin |
|---|---|---|---|
| H3-GFP | H3.1-EGFP | pEGFP-N1 | Apta-Smith et al., 2018. |
| H3 HB-GFP | H3.1 (A95_GGG)-EGFP | pEGFP-N1 | This study |
| H3 FD-GFP | H3.1 (FLY>AAA)-EGFP | pEGFP-N1 | This study |
| H4-GFP | H4-EGFP | pEGFP-N1 | Apta-Smith et al., 2018. |
| H4 HB-GFP | H4 (V65GGG)-EGFP | pEGFP-N1 | This study |
| H4 FD-GFP | H4 (FLI>AAA)-EGFP | pEGFP-N1 | This study |
| H3-mCherry-RR | H3.2-mCherry-2xTVMVcs-FRB-OMP5–IRES–FKBP-TVMV-AI | pEGFP-C1 | This study |
| GFP-Imp5 | EGFP-Imp5 | pEGFP-C1 | This study |
| GFP-sNASP | EGFP-TEV-sNASP | pIRESpuro2 | This study |
| H3-APEX2-EGFP-RR | H3.1-Flag-APEX2-EGFP-2xTVMVcs-FRB-OMP25-IRES-FKBP-TVMV-AI | pcDNA™5/FRT/TO | This study |
| NASP_HDR | pNASP-EGFP-TEV_HDR-donor | pBlueScript II KS (+) | This study |
| NASP_sgRNA_Down | pX461-NASP-sgRNA2-down | pX461-PSPCas9N(BB)-2A-GFP | This study |
| NASP_sgRNA_up | pX461-NASP-sgRNA2-up | pX461-PSPCas9N(BB)-2A-GFP | This study |
| ASF1B_HDR | pASF1b-Spot-mCherry-3C_HDR-donor | pBlueScript II KS (+) | This study |
| ASF1B_sgRNA_Down | pX461-ASF1B-sgRNA2-down | pX461-PSPCas9N(BB)-2A-GFP | This study |
| ASF1B_sgRNA_up | pX461-ASF1B-sgRNA2-up | pX461-PSPCas9N(BB)-2A-GFP | This study |
| GST-Imp5 | pGST-HRV 3Ccs-Imp5 iso1 | pGEX-6P1 | This study |
