## Supplementary material for "A specific role for Importin-5 and NASP in the import and nuclear hand-off of monomeric H3": Table S2

**Table S2. Primers and gene fragments sequences**

| Primer_name | Sequence |
|---|---|
| XhoI_IPO5_F | aaaaaaCTCGAGcaATGGCGGCGGCCGC |
| KpnI_IPO5_R | GGTGGTggtaccTCACGCAGAGTTCAGGAGCTC |
| IPO5_pGEX_F | ctgttccaggggcccctgggatccGCAATGGCGGCGGCtGCG |
| IPO5_pGEX_R | gtcagtcagtcacgatgcggccgcTCACGCAGAGTTCAGGAGC |
| Kozak5p_F | TGAACCGTCAGATCCGCTAGC |
| OMP25_IRES_R | CGGTAGCGCTACAGCTGTTTGCGATAGCG |
| OMP25_IRES_F | TCGCAAACAGCTGTAGCGCTACCGGACTCAG |
| IRES_FKBP_R | TCCACCTGCACCATGGTTGTGGCCATAT |
| FKBP_IRES_F | ATGGCCACAACCATGGTGCAGGTGGAAACC |
| FKBP_FRT_R | TGTGGGAGGTTTCTAGCTGCCCGGCGC |
| FRT_AI_F | CCGGGCAGCTAGAAACCTCCCACACCTCCCCCT |
| Kozak_H3.1_R | GATCTGACGGTTCACTAAAC |
| APEX2_gBLOCK | AGCGCTACCGGTCGCCACCATGGACTACAAGGATGACGACGATAAGGGAAAGTCTTACCCAACTGTGAGTGCTGATTACCAGGACGCCGTTGAGAAGGCGAAGAAGAAGCTC<br>AGAGGCTTCATCGCTGAGAAGAGATGCGCTCCTCTAATGCTCCGTTTGGCATTCCACTCTGCTGGAACCTTTGACAAGGGCACGAAGACCGGaGGACCCTTCGGAACCATCAA<br>GCACCCTGCCGAACTGGCTCACAGCGCTAACAACGGTCTTGACATCGCTGTTAGGCTTTTGGAGCCACTCAAGGCGGAGTTCCCTATTTTGAGCTACGCCGATTTCTACCAGTT<br>GGCTGGCGTTGTTGCCGTTGAGGTCACGGGTGGACCTAAGGTTCCATTCCACCCTGGAAGAGAGGACAAGCCTGAGCCACCACCAGAGGGTCGCTTGCCCGATCCCACTAAG<br>GGTTCTGACCATTTGAGAGATGTGTTTGGCAAAGCTATGGGGCTTACTGACCAAGATATCGTTGCTCTATCTGGGGGTCACACTATTGGAGCTGCACACAAGGAGCGTTCTGGA<br>TTTGAGGGTCCCTGGACCTCTAATCCTCTTATTTTCGACAACTCATACTTCACGGAGTTGTTGAGTGGTGAGAAGGAAGGaCTCCTTCAGCTACCTTCTGACAAGGCTCTTTTGT<br>CTGACCCTGTATTCCGCCCTCTCGTTGACAAATATGCAGCGGACGAAGATGCCTTCTTTGCTGATTACGCTGAGGCTCACCAAAAGCTcTCCGAGCTTGGGTTTGCTGATGCCgg<br>taccggaggtAAGTACTCAGATCTCGAGggaggttccggcGAGCTCAAGCTTCGCGGCGATGGCGAACCGAGCGGCGTGCCGGTGGCGGTGGTGCTGCTGCCGGTGTTTGCGCTGACC<br>CTGGTGGCGGTGTGGGCGTTTGTGCGCTATCGCAAACAGCTGTAGGGATCCACCGGATCTAGATAACTGATCATAATCAGCCATAC |
| NASP_HDR_upstream_F | AGCTACTCGCCCTGAACATGCAGAGCAGCACTG |
| NASP_HDR_upstream_R | CGTTCCCCTGAGGTGGCGAACCAGCGAACG |
| GFPforNASP_HDR_F | TTCGCCACCTCAGGGGAACGATGGTGAGCAAGGGCGAGGA |
| GFPforNASP_HDR_R | GCTGTGGACTCCATGGCCATATGGCCCTGGAAGTAAAGGT |

| | |
|---|---|
| NASP_HDR_downstream_F | ATGGCCATGGAGTCCACAGCCACTGCCGC |
| NASP_HDR_downstream_R | TTATATTCCCGGGTTATCCAGGGGTTCTACCAGAGGCACACG |
| NASP_pair1_upstream_F | CACCGCCATGGCCATCGTTCCCCTG |
| NASP_pair1_upstream_R | AAACCAGGGGAACGATGGCCATGGC |
| NASP_pair1_downstream_F | CACCGAGCCACTGCCGCCGTCGCCG |
| NASP_pair1_downstream_R | AAACCGGCGACGGCGGCAGTGGCTC |
| ASF1B_UP_R | GGCCATCGCCTCGCCTCGCC |
| ASF1B_UP_F | AATCACTTCGGGTGCGAGCACC |
| Spot-tag_mCherry_3C_R | GCACCGACACCTTGGCCATGGGCCCCTGGAACAGAACTTCCAGGAGTCCGGACTTGTACAGCT |
| Spot-tag_mCherry_3C_F | GAGGCGAGGCGatggccccggatcgcgtgcgcgcggtgagccattggagcagcGTGAGCAAGGGCGAGGAGGA |
| ASF1B_DOWN_R | GGAAAATGGGAAGGGGCTGGATATTGG |
| ASF1B_DOWN_F | ATGGCCAAGGTGTCGGTGC |
| ASF1B_sgRNA_UP_F | CACCGTCGCCTCGCCGCGCCGCAGC |
| ASF1B_sgRNA_UP_R | AAACGCTGCGGCGCGGCGAGGCGAC |
| ASF1B_sgRNA_Down_F | CACCGCAAGGTGTCGGTGCTGAACG |
| ASF1B_sgRNA_Down_R | AAACCGTTCAGCACCGACACCTTGC |
