## Supplementary material for "A specific role for Importin-5 and NASP in the import and nuclear hand-off of monomeric H3": Table S3

**Table S3. Antibodies, companies, catalogue number and lot, dilutions used and webpage link**

| Name | Company | Catalogue number | Lot | Origin | Concentration used | Webpage |
|---|---|---|---|---|---|---|
| Anti-ASF1A (C6E10) | Cell Signaling Technology | C6E10 | | Rabbit - monoclonal | 1:1 k | https://media.cellsignal.com/pdf/2990.pdf |
| Anti-ASF1B | Cell Signaling Technology | C70E2 | 2 | Rabbit - monoclonal | 1:1 k | https://www.cellsignal.co.uk/products/primary-antibodies/asf1b-c70e2-rabbit-mab/2902 |
| Anti-HAT1 | Abcam | ab194296 | GR239204-4 | Rabbit - monoclonal [EPR18775] to KAT1 / HAT1 | 1:1 k | https://www.abcam.com/kat1--hat1-antibody-epr18775-ab194296.html |
| Anti-Histone H3 | Cell Signaling Technology | #9715 | 23 | Rabbit - polyclonal, IgG | 1:1 k | https://www.cellsignal.co.uk/products/primary-antibodies/histone-h3-antibody/9715 |
| Anti-Histone H4 | Abcam | ab7311 | GR274242-1 | Rabbit - polyclonal to Histone H4 - ChIP Grade, IgG | 1:1 k | https://www.abcam.com/histone-h4-antibody-chip-grade-ab7311.html |
| Anti-Histone H4 k12Ac | Millipore (Merck) | #07-595 | #3328962 | Rabbit - polyclonal | 1:1 k | https://www.merckmillipore.com/GB/en/product/Anti-acetyl-Histone-H4-Lys12-Antibody,MM_NF-07-595 |
| Anti-Histone H4 k5Ac | Millipore (Merck) | #07-327 | #3307312 | Rabbit - polyclonal | 1:1 – 5 k | https://www.merckmillipore.com/GB/en/product/Anti-acetyl-Histone-H4-Lys5-Antibody,MM_NF-07-327?ReferrerURL=https%3A%2F%2Fwv |
| Anti-Importin4 | Abcam | ab181037 | GR151286-1 | Rabbit - monoclonal [EPR13660-27] to Importin4 | 1:10 k | https://www.abcam.com/importin4-antibody-epr13660-27-ab181037.html |
| Anti-Karyopherin beta 3 (IPO5) | Abcam | ab88695 | GR3031337-1 | Mouse polyclonal | 1:500 50 µg at 1 mg/ml, secondary at 1: 5 k | https://www.abcam.com/karyopherin-beta-3-antibody-ab88695.html |
| Anti-mCherry | Abcam | ab167453 | GR3265215-1 | Rabbit polyclonal | 1:5 k | https://www.abcam.com/mcherry-antibody-ab167453.html?productWallTab=ShowAll |
| Anti-RbAp46/48 | Cell Signaling Technology | (D4F8) #9067 | 1 Ref 10/2017 | Rabbit - monoclonal (IgG) | 1:1 k | https://www.cellsignal.co.uk/products/primary-antibodies/rbap46-rbap48-d4f8-rabbit-mab/9067?Ntk=Products&Ntt=9067 |
| Anti-UBR7 | Abcam | ab241371 | GR3262878-3 | Rabbit - polyclonal | 1:1 k | https://www.abcam.com/ubr7-antibody-ab241371.html |
| GFP (B-2) HRP | Santa Cruz | sc-9996 HRP | I2618 | Mouse - monoclonal (clone B-2), IgG2a | 1:1 to 1:15 k | https://datasheets.scbt.com/sc-9996.pdf |
| Goat Anti-Mouse HRP | Abcam | ab205719 | GR247077-9 | Goat - polyclonal, IgG | 1:5 k | https://www.abcam.com/goat-mouse-igg-hl-hrp-ab205719.html |
| Anti-Rabbit, HRP-linked antibody | Cell Signaling Technology | 7074S | 27 | Goat - polyclonal, IgG | 1:10 – 20 k | https://www.cellsignal.co.uk/products/secondary-antibodies/anti-rabbit-igg-hrp-linked-antibody/7074 |
| Goat Anti-Rabbit Alexa 568 | Abcam | ab175471 | GR1811195-3 | Goat - polyclonal, IgG | 1:1 k | https://www.abcam.com/goat-rabbit-igg-hl-alexa-fluor-568-ab175471.html |
